# An Altered Glycome Shapes IgA B-Cell Responses and Gut Immunity During Intestinal Inflammation

**DOI:** 10.64898/2026.06.08.730456

**Authors:** Anabela M. Cutine, Alejandro J. Cagnoni, Joaquín P. Merlo, Ayelén D. Rosso, Pablo A. García, Montana N. Manselle Cocco, Luciano G. Morosi, Mora Massaro, Verónica C. Martínez Allo, Rosa M. Morales, Sabrina G. Gatto, María May, Diego O. Croci, Renata Spiazzi, Carolina Conlon, Jimena Cerezo, Claudia Milano, Alberto Penas-Steinhardt, Fiorella S. Belforte, Gabriel A. Rabinovich, Marta A. Toscano, Karina V. Mariño

## Abstract

A healthy gut immune system balances pathogen defense and tolerance to beneficial microbes. This equilibrium is sustained by coordinated mechanisms where B cells (BCs) play a central role, and secretory immunoglobulin A (SIgA) regulates microbiome composition. In ulcerative colitis (UC), impaired tolerogenic pathways result in exaggerated immune activation, epithelial dysfunction, and tissue damage; however, the contribution of BCs to disease pathogenesis remains unclear. Notably, sialylation is crucial to B-cell function, but its relevance in the intestinal IgA B-cell response has been scarcely explored.

Here, we show that SIgA from active UC patients displays an inflammation-dependent reduction in α(2,6)-sialylation. This desiaylation is recapitulated in dextran sodium sulfate–induced colitis, where IgA⁺ plasma cells (IgA^+^ PCs) and BCs exhibit a similar glycophenotype. Functional analyses reveal that BCs lacking α(2,6)-sialylation on N-glycans exhibit defective differentiation into IgA⁺ PCs and diminished capacity to suppress intestinal inflammation *in vivo*, with increased neutrophil infiltration. Moreover, transcriptomic analyses of UC patient samples suggest a synergistic contribution of neuraminidase activity and reduced bioavailability of sialic acid precursors, leading to SIgA desialylation. Collectively, these findings uncover a glycosylation-dependent pathological circuit in which altered sialylation of SIgA and BCs disrupts their function, compromising mucosal homeostasis in intestinal inflammation.

## INTRODUCTION

With an estimated prevalence of 5 million cases worldwide, ulcerative colitis (UC) greatly impacts quality of life, leading to serious health complications including increased risk of colorectal cancer [1,2]. Biological therapies such as anti-tumor necrosis factor, anti-integrin α_4_β_7_, and more recently, anti-IL-12, anti-IL-23 and Janus kinase inhibitors have significantly improved treatment options; however, remission rates still range around 30%-60% of patients, and medically-refractory disease still leads to surgical intervention [1]. Epidemiological studies indicate that the global burden of inflammatory bowel diseases (IBD), including UC, will continue to rise with a rapid increase in incidence in newly industrialized countries and regions of Asia and Latin America [3].

The most widely accepted etiopathogenic mechanism links UC to defective immune tolerance pathways, leading to exacerbated responses to intestinal microbiota, epithelial dysfunction, and tissue damage [1]. While most experimental and clinical studies in UC have focused on T cells, macrophages, and soluble cytokines such as TNF and IL-23, basal plasmacytosis (a dense infiltration of plasma cells) is a typical feature of UC and has been proposed as an indicator of severity or relapse [4]. The presence of autoantibodies and dysbiosis in UC patients brought B cells, and in particular, plasma cells (PCs) back to the spotlight [5–9]. Notably, experimental models have shown both suppressive [10–12] and detrimental roles [6] for B cells in intestinal inflammation, highlighting their multifaceted roles. Clinically, recent evidence repositions therapeutic B-cell targeting in UC, as CD19 CAR T-cell therapy has led to drug-free remission and mucosal healing [13].

Secretory IgA (SIgA) is the primary immunoglobulin in the healthy gut, providing the first line of immune defense against pathogens and shaping intestinal microbiota through mechanisms that include neutralizing bacterial toxins and antigens [14,15]. In SIgA, IgA dimers are covalently bound by the J chain and interact with the secretory component (SC), which is derived from epithelial cells during transcytosis [14]. Notably, in UC patients increased intestinal IgG production and induction of anti-commensal IgG correlated with disease severity [16–18]. Moreover, the increase in hyperactive IgG^+^ PCs has been proposed to participate in the pathogenesis of this pathology [7,8,18]. However, IgA^+^ PCs remain an important population within the gut PC-compartment in UC, and the potential functional alterations of SIgA in this disease are only beginning to be explored [14,19].

Glycans and glycan-binding proteins have emerged as key players in the regulation of intestinal immune homeostasis, particularly in the context of pathogen recognition, immune cell activation and trafficking [20–24]. In addition, extensive work demonstrated the relevance of sialylation in B cells [25,26] and IgG function [27,28]. In autoimmune disorders, agalactosylated and desialylated N-glycans confer proinflammatory activity to serum IgG, and this altered IgG glycome has been proposed as a prognostic parameter for monitoring development and progression of rheumatoid arthritis [27], and correlates with disease severity in IBD [29].

IgA and SIgA are extensively glycosylated; however, research focusing on their glycosylation and its functions remains limited. Studies on serum IgA revealed that IgA2 glycans show a more immature phenotype, with a higher prevalence of oligomannose glycans and fewer sialylated N-glycans than IgA1 [30,31]. Functionally, IgA2 exhibits proinflammatory effects on macrophages and neutrophils, while desialylation or de-N-glycosylation of IgA1 increases its inflammatory activity, highlighting glycans as modulators of IgA effector functions [31]. Recently, serum IgA glycosylation in Crohńs disease has been described, with lower N-glycan sialylation [32]. Notably, α(2,6)-sialylation has been reported as the dominant sialic acid linkage in SIgA N-glycans [33] and glycosylation of the SC safeguards SIgA from proteolytic degradation and facilitates its attachment to mucus [15]. Additionally, glycosylation of the J chain and SC is implicated in alternative antigen binding through non-Fab sites (non-canonical binding) [15]. Polyreactive IgA, which is induced by commensal microbes and dietary antigens, also engages in non-specific glycan-mediated interactions [14]. Finally, recent studies proposed that in UC SIgA loses its effector functions, displaying low levels of reverse transcytosis and anti-glycan reactivity [19].

In this work, we demonstrate that gut SIgA α(2,6)-sialylation is decreased during intestinal inflammation, both in UC patients and in the DSS-induced colitis model. Furthermore, we show that intestinal B cells and IgA^+^ PCs in experimental colitis also exhibit this altered sialylation. Focusing on the functional consequences of this aberrant glycosylation profile, we show that B cells devoid of α(2,6)-sialic acid have a defective capacity to suppress inflammation and an impaired differentiation to IgA-producing PCs. Based on these findings we explored the potential causes of this altered sialylation profile, including dysregulation of the glycosylation machinery in IgA^+^ PCs and epithelial cells, proposing mechanisms that could mediate aberrant SIgA glycosylation in UC.

## RESULTS

### SIgA α(2,6)-sialylation is decreased in active UC, but not in remission

To determine if sialylation of secretory IgA (SIgA) is altered in ulcerative colitis (UC), we performed lectin blotting with SIgA isolated from fecal samples of UC patients and non-IBD individuals (**Fig. 1A**, **Table S1**). *Sambucus nigra* agglutinin (SNA, with affinity for α(2,6)-sialic acid, **Fig. S1A**), demonstrated a significant decrease in α(2,6)-sialylation in the heavy chain (HC) of fecal SIgA from active UC patients (**Fig. 1B,C**), with a similar trend in the SC (**Fig. 1B,D**). Given that desialylation exposes galactose residues on N-glycans, we evaluated SIgA galactosylation by *Erythrina cristagalli* lectin (ECL) binding (**Fig. S1A**). Results showed increased exposure of galactose residues in both HC and SC from SIgA in active UC (**Fig. 1B-D**), although no significant differences were observed in sialylation or galactosylation profiles between UC patients in remission and non-IBD individuals (**Fig 1E-G**). Hypothesizing that the loss of α(2,6)-sialic acid would alter SIgA functionality, we then evaluated bacterial binding. As a proof of concept, we removed sialic acid from human SIgA (hSIgA) and measured its binding to fecal bacteria from *Rag2*^-/-^ mice, which lack endogenous immunoglobulins (**Fig. S1B**). Interestingly, desialylated hSIgA shows increased binding to *Rag2*^-/-^ fecal bacteria (**Fig. S1C,D**). Thus, SIgA shows aberrant glycosylation in active UC, a trait that could affect its functionality.

**Figure 1.**
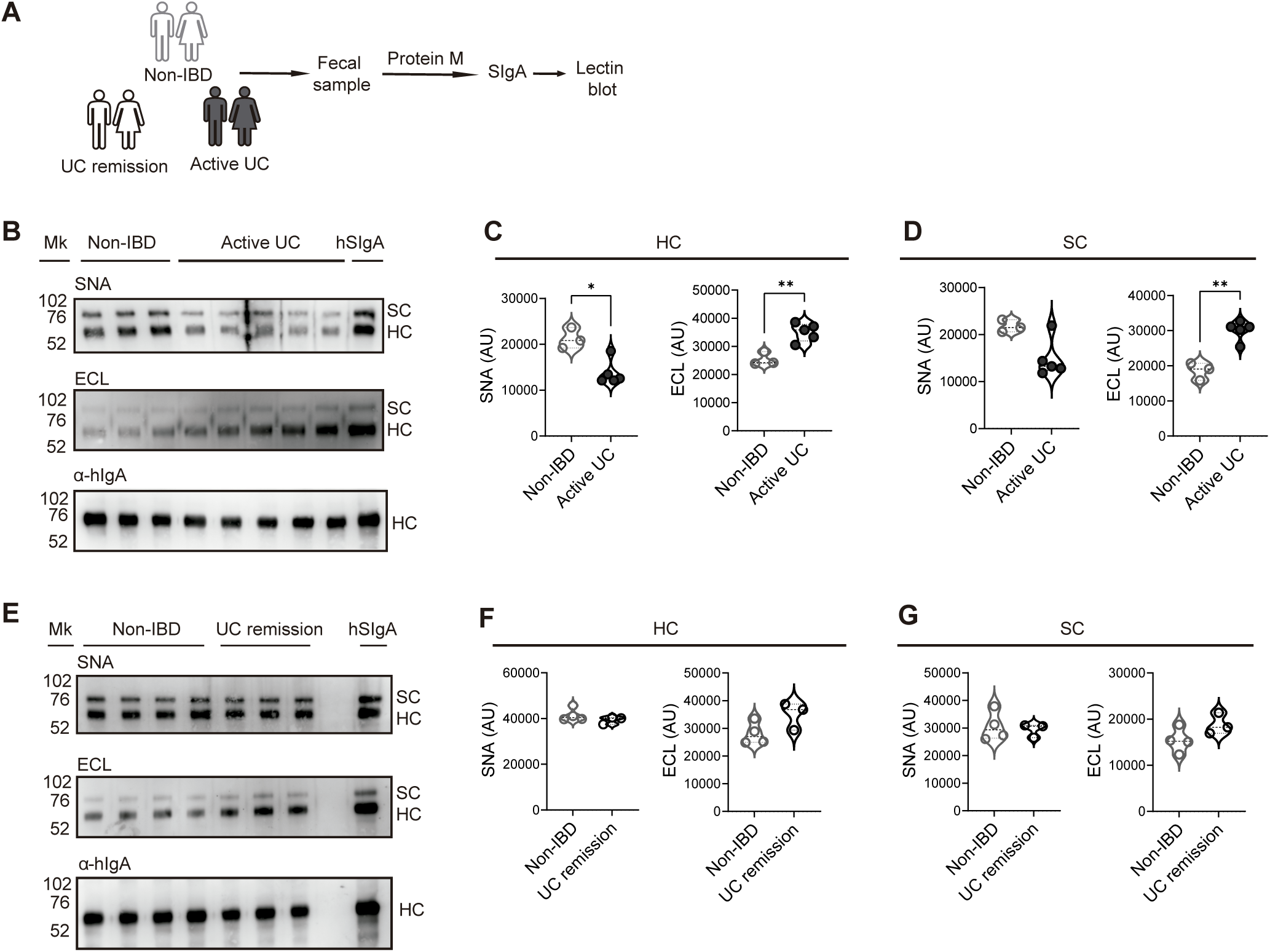
Fecal SIgA is aberrantly sialylated in active UC. **(A)** Study design and SIgA purification. **(B)** α(2,6) sialylation and galactosylation measured by SNA and ECL blotting, respectively, in SIgA samples isolated from non-IBD individuals and active UC patients. A western blot with anti-human IgA is shown as a reference and loading control, and human colostrum SIgA (hSIgA) is used as a control. **(C,D)** SNA and ECL binding to the SIgA heavy chain (**C,** HC, 55 kDa) and secretory component (**D,** SC, 75 kDa) (violin plots, each dot represents one patient, quartiles and median shown as dotted lines; n = 3-5). **(E)** SNA and ECL lectin blots performed with SIgA isolated from non-IBD individuals and UC patients in remission. **(F, G)** SNA and ECL binding to the HC **(F)** and SC **(G)** of SIgA from non-IBD individuals and UC patients in remission (violin plots, each dot represents one patient, quartiles and median shown as dotted lines; n = 3-4). Student’s t test for all comparisons except SNA binding to HC and SC (Mann Whitney), *p<0.05. AU, arbitrary units.

### α(2,6)-sialylation on SIgA, IgA^+^ plasma cells and B cells is decreased in DSS-induced colitis

To understand the mechanisms underlying SIgA aberrant sialylation, we conducted experiments in a chronic DSS-colitis model [22](**Fig. 2A,B**). DSS-treated mice showed increased IgA in colon and serum (**Fig. 2C,D**), which negatively correlated with colonic length (**Fig. 2E**, **Fig. S2A**), indicating an association with disease severity. Remarkably, intestinal SIgA from DSS-treated mice showed a significant reduction in α(2,6)-sialic acid, recapitulating the profile observed in SIgA from active UC patients (**Fig. 2F**). We hypothesized that this aberrant SIgA glycoprofile could be mirrored by IgA^+^ PCs and consequently analyzed the glycophenotype of colonic lamina propria (LP) IgA^+^ PCs (**Fig. 2G**). Interestingly, both confocal microscopy (**Fig. S2B**) and flow cytometry showed a significant increase in LP CD138^+^IgA^+^ PCs in DSS-colitis (**Fig. 2G**, **Fig. S2C**). IgA^+^ PCs from control mice showed cell surface α(2,3)- and α(2,6)-sialylation, as shown by *Maackia amurensis* lectin II (MALII) and SNA binding, respectively, with very low abundance of β(1,6)-branched N-glycans (as determined by PHA-L binding), polylactosamine extensions by *Lycopersicon esculentum* lectin (LEL), and asialo-core-1 O-glycans (PNA) (**Fig. 2H**, **top panel**). Notably, LP IgA^+^ PCs from DSS-treated mice showed reduced α(2,3)- and α(2,6)-sialylation when compared to controls, with no significant differences in other lectins tested (**Fig. 2H**). Thus, LP IgA^+^ PCs recapitulate the aberrant sialylation profile detected in SIgA.

**Figure 2.**
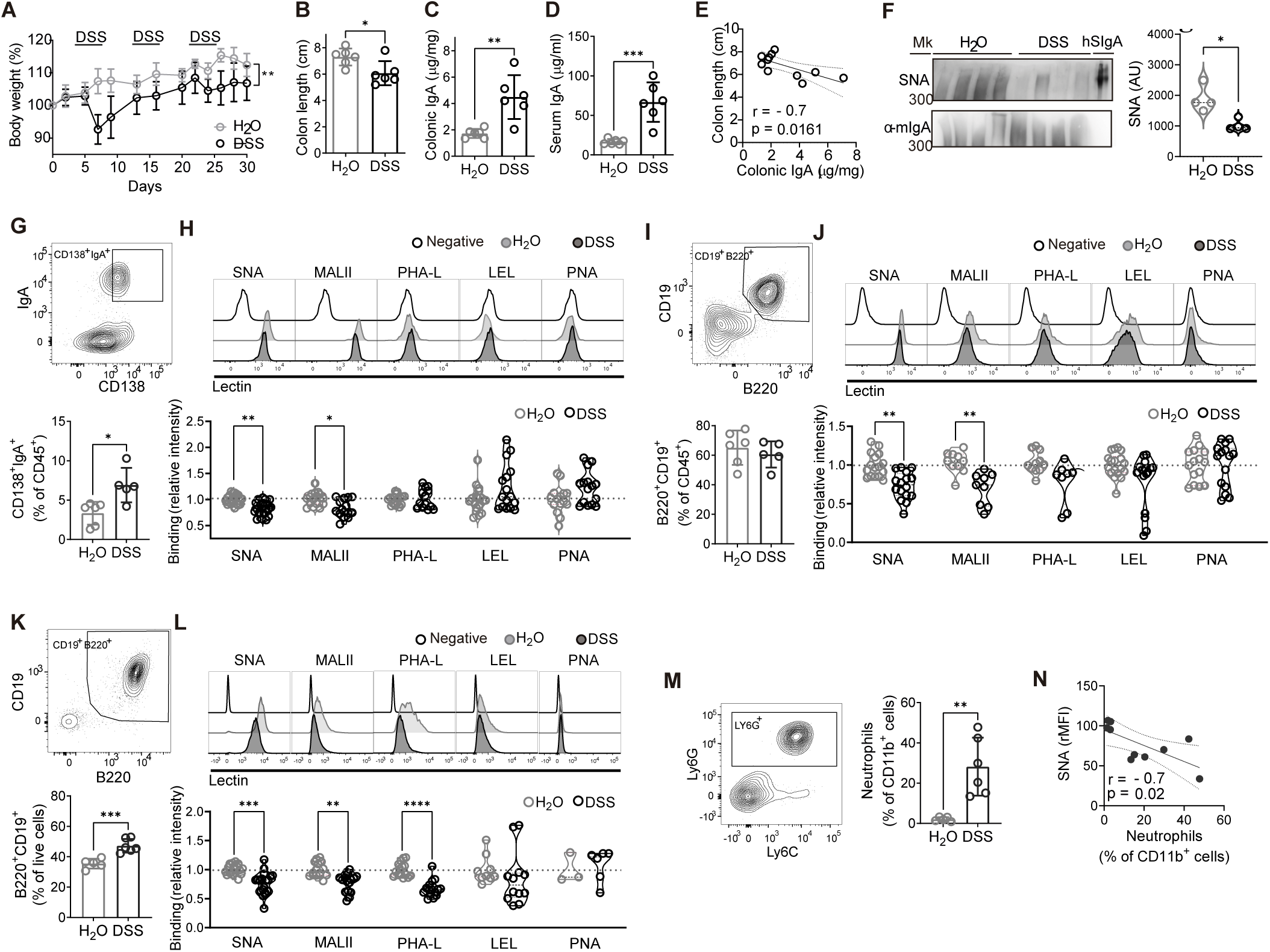
Sialylation of SIgA, IgA^+^ PCs and B cells is altered in DSS-induced colitis. **(A)** Weight loss in chronic DSS colitis (DSS) and control mice (H_2_O) (mean ± SD; n = 6, data representative of 3 independent experiments; two-way repeated-measures ANOVA followed by Sidak’s test). **(B)** Colon length (mean ± SD; n = 6; Student’s t-test). **(C,D)** Colonic and serum IgA by ELISA (mean ± SD, n = 6; Student’s t-test). **(E)** Correlation between colon length and colonic IgA in control and DSS-treated mice (n = 11). **(F) Left:** Lectin-blot (SNA) and western-blot (anti-mouse IgA) of luminal SIgA in non-reducing conditions. hSIgA is shown as control. **Right:** SNA binding to mouse IgA (violin plots, n = 4, each sample is pooled from 2-4 mice from two independent experiments, Mann-Whitney test). AU, arbitrary units. **(G-L)** Proportion and glycophenotype of LP CD138^+^IgA^+^ PCs **(G,H),** LP B cells **(I,J)** and MLN B cells **(K,L). (G, I, K)** Dot plot and proportions of LP IgA^+^ PCs (**G)**, LP B cells (**I**) and MLN B cells (**K**) (mean ± SD; n = 5-6, data representative of three or four independent experiments, Student’s t test for all comparisons except IgA^+^ PCs (Mann-Whitney test). **(H, J, L) Top:** Representative histograms of biotinylated lectin binding to LP CD138^+^IgA^+^ PCs **(H)**, LP B cells **(J)** and MLN B cells **(L)**; **Bottom**: Relative intensity of lectin binding. For (**H**) and (**J**), violin plots; n = 8-22, data pooled from two to four independent experiments; Kruskal–Wallis test followed by Dunn’s multiple comparisons test. For (**L**), violin plots; n = 3-19, data pooled from one to four independent experiments; Kruskal–Wallis test followed by Dunn’s multiple comparisons test. (**M**) Dot plot for LP neutrophils and proportion of these cells (mean ± SD; n = 5-6, Student’s t test). **(N)** Correlation between LP neutrophils and SNA binding to MLN B cells (n=11). For (**D,N**), solid black line indicates the best fit curve, dotted lines indicate 95% trust intervals. Pearsońs P-value (p) and correlation coefficient (r) are shown, *p<0.05, **p<0.01, ***p<0.0001.

Of note, when we examined LP lymphocytes in DSS-colitis (gating strategy in **Fig. S2C**), we found no significant differences in the proportions of B cells (**Fig. 2I**) or CD8^+^ or CD4^+^ T cells (**Fig. S2D**). Similarly to IgA^+^ PCs, LP B cells in control mice showed high levels of cell surface α(2,6)- and α(2,3)-sialic acid with low PNA binding; however, these cells showed considerable levels of β(1,6)-branched N-glycans and poly-LacNAc extensions (**Fig. 2J**, **top panel**). Remarkably, in DSS-treated mice, LP B cells also show a reduction in α(2,3)- and α(2,6)-sialylation with no differences in other lectins when compared with controls (**Fig. 2J**). Notably, we found a significant increase in the proportion of B cells in mesenteric lymph nodes (MLN) during colitis (**Fig. 2K**). Similarly to LP B cells, MLN B cells of control mice showed high levels of α(2,6)- and α(2,3)-sialic acid, β(1,6)-branched N-glycans and polylactosamine extensions, and very low levels of asialo-core-1 O-glycans (**Fig. 2L**, **top panel**). Interestingly, MLN B cells mirrored LP IgA^+^ PCs and B cells in DSS-colitis, with decreased cell surface α(2,3)- and α(2,6)-sialylation; however, β(1,6)-branched N-glycans were also reduced, with no other significant differences (**Fig. 2L**). Hence, intestinal inflammation modulates sialylation not only in LP IgA^+^ PCs, but also in MLN and LP B cells.

Given the key contribution of innate immunity to intestinal inflammation [34], myeloid cells were evaluated by flow cytometry. Chronic DSS treatment led to a significantly higher proportion of neutrophils when compared with controls (**Fig. 2M**). Furthermore, we also observed higher proportions of intermediate monocytes (Ly6C^hi^MHCII^hi^) and decreased frequency of mature macrophages (Ly6C^low^MHCII^hi^), with no alterations in LP CD11b^+^ total myeloid cells (**Fig. S2E,F**). Notably, we detected a negative correlation between α(2,6) sialic acid in MLN B cells and the proportion of LP neutrophils (**Fig. 2N**), indicating a potential link between inflammation and B-cell sialylation.

In summary, gut SIgA α(2,6)-sialylation decreases in DSS-induced colitis, as in fecal SIgA from UC patients, and this aberrant sialylation profile is recapitulated in IgA^+^ PCs and B cells in this experimental model.

### Genetic ablation of α(2,6)-sialylation in B-cell N-glycans exacerbates intestinal inflammation

The observed decrease in α(2,6)-sialylation within SIgA, IgA^+^ PCs, and B cells during DSS-induced inflammation prompted us to investigate the relevance of α(2,6)-sialylation in B cell functionality during gut inflammation. Considering that β-galactoside α(2,6)-sialyltransferase 1 (ST6GAL1) is a highly conserved enzyme responsible for attaching α(2,6)-sialic acid to N-glycans [25], we generated transgenic mice with a B cell-specific deletion of *St6gal1* (*Cd19*^Cre^ *St6gal1*^f/f^; St6 B-KO). Validation of the phenotype by SNA binding showed selective deletion of α(2,6)-sialic acid in the B cell-compartment from St6 B-KO mice and normal sialylation in CD4^+^ T cells (**Fig. S3A**).

Notably, DSS-induced colitis in St6 B-KO mice resulted in greater colonic shortening compared to control littermates (*St6gal1*^f/f^; St6 flox), although weight loss was not significantly different (**Fig. 3A,B**). Additionally, St6 B-KO mice with colitis exhibited a decreased frequency of B cells and IgA^+^ PCs (**Fig. 3C**) with no significant changes in T cell populations (**Fig. 3D**). Notably, these mice also showed a significant increase in CD11b^+^ myeloid cells, mainly neutrophils and eosinophils, and similar trends in monocytes and dendritic cells (**Fig. 3E,F**, **Fig. S3B**). Moreover, neutrophil infiltration correlated with colonic shortening (**Fig. S3C**). In line with these findings, St6 B-KO mice show a trend toward higher inflammatory markers, including serum lipocalin (**Fig. 3G**), colonic IL-1α, IL-1β, and IL-17A (**Fig. S3D**).

**Figure 3.**
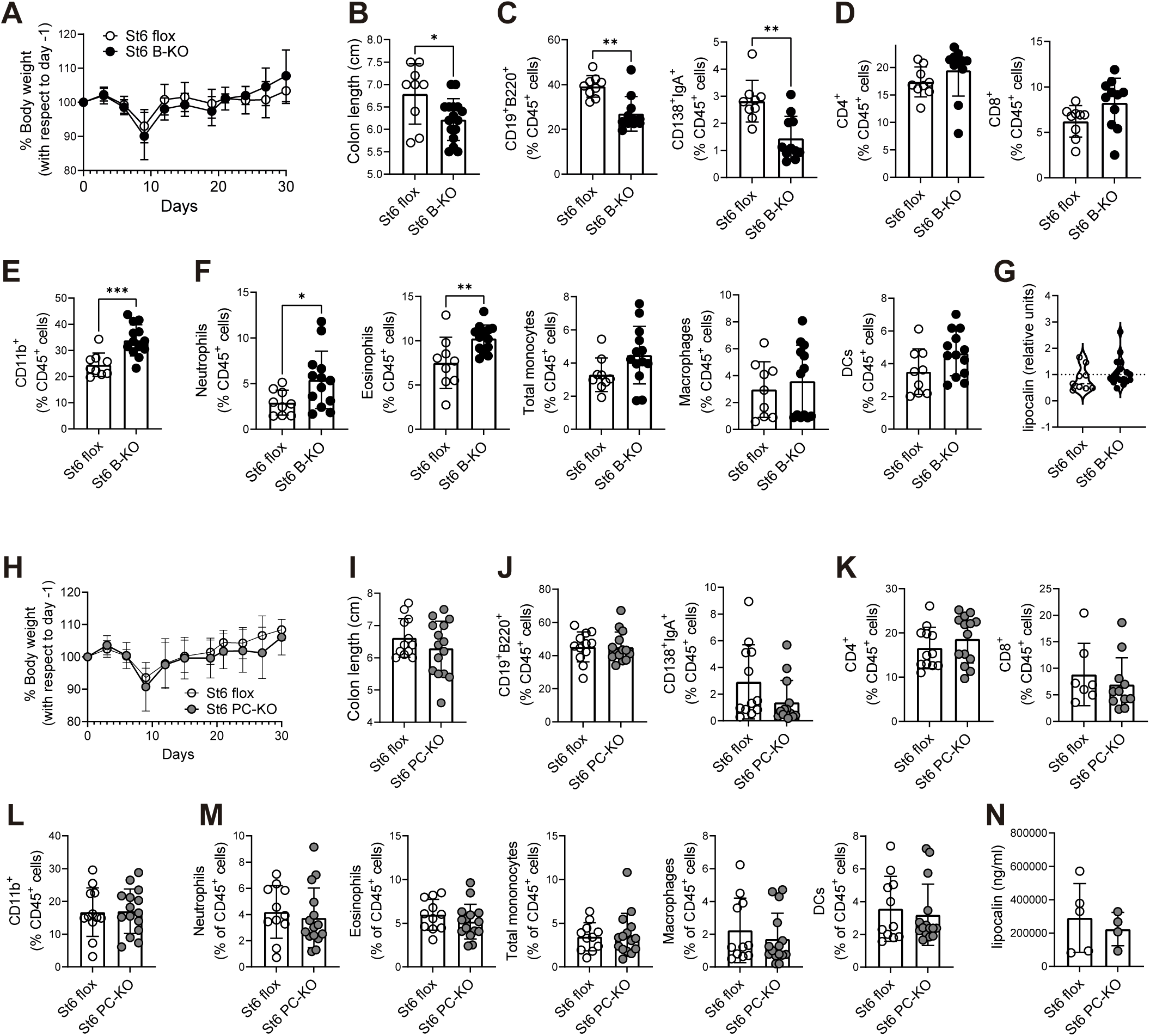
B-cell specific *St6gal1* deficiency aggravates DSS-induced colitis. **(A-G)** DSS-induced colitis in *Cd19^Cre^ St6gal1*^f/f^ (St6 B-KO) and *St6gal1*^f/f^ (St6 flox) *littermates*. **(A)** Weight loss curve (mean ± SD; n = 4-7, data representative of two independent experiments). **(B)** Colon length. **(C-F)** Proportions of LP immune cells. **(C)** Proportions of B cells and CD138^+^IgA^+^ PCs, **(D)** CD4^+^ and CD8^+^ T cells and **(E)** CD11b^+^ total myeloid cells and **(F)** myeloid subpopulations. **(B-F)**: mean ± SD; n = 9-17, data from two independent experiments; Student’s t test for all comparisons except B cells, CD4^+^ cells and macrophages (Mann-Whitney test). **(G)** Violin plot depicting relative levels of serum lipocalin by ELISA (n = 9-17, data are from two independent experiments, levels of lipocalin within each experiment were relativized by dividing with corresponding experiment mean; Mann Whitney test). **(H-N)** DSS-induced colitis in *Aicda^Cre^ St6gal1*^f/f^ (St6 PC-KO) and *St6gal1*^f/f^ (St6 flox) (St6 PC-KO) littermate mice. **(H)** Weight loss curve (mean ± SD; n = 3-6, data representative of three independent experiments). **(I)** Colon length. **(J-M)** Proportions of LP immune cells. **(J)** Proportions of LP B cells, CD138^+^IgA^+^ PCs, **(K)** CD4^+^ and CD8^+^ T cells and **(L)** CD11b^+^ total myeloid cells and **(M)** myeloid subpopulations. **(I-M)** mean ± SD; n = 12-15, data from three independent experiments, Student’s t test. **(N)** Levels of serum lipocalin measured by ELISA (mean ± SD; n = 4-5, data from one experiment; Student’s t test). *p<0.05; **p<0.01; ***p<0.001; ****p<0,0001.

Considering that in St6 B-KO mice B cells are devoid of α(2,6)-sialic acid in all developmental stages, we next investigated the functionality of α(2,6)-sialylation specifically in PCs. Thus, we generated mice with PCs deficient in *St6gal1* (*Aicda*^Cre^ *St6gal1*^f/f^; St6 PC-KO). Comparative SNA binding of IgA^+^ PCs and CD4^+^ T cells confirmed the loss of α(2,6)-sialic acid specifically in the former (**Fig. S3E**). Interestingly, St6 PC-KO mice showed a trend towards a decrease in colon IgA^+^ PCs (**Fig. 3J**), with no differences in colon length (**Fig. 3I**), weight loss (**Fig. 3H**), frequency of LP immune cells (**Fig. 3J-M**) or inflammatory markers (**Fig. 3N**, **Fig. S3F**) when compared to littermate controls (*St6gal1*^f/f^; St6 flox).

Collectively, these results indicate that lack of α(2,6)-sialic acid in B cells alters the immune infiltrate, with decreased frequency of B cells and IgA^+^ PCs and increased innate immune infiltration and inflammation in the colon. Conversely, deletion of α(2,6)-sialic acid exclusively on PCs does not recapitulate this phenotype.

### B cells lacking α(2,6)-sialylation showed impaired differentiation to PCs and decreased suppressive function *in vivo*

Based on our findings, B cells deficient in α(2,6)-sialic acid play a pathogenic role during DSS-colitis. To further investigate the mechanisms involved and rule out model-specific artifacts, we functionally assessed these cells in the T-cell transfer colitis model, where the suppressive role of B cells has been previously established [12]. For this purpose, we transferred WT CD4^+^CD45RB^hi^ colitogenic naive T cells into *Rag2*^-/-^ mice and co-transferred either *St6gal1*^-/-^ (St6-KO) or WT B cells (**Fig. 4A**). As expected, WT B cells partially suppressed T cell-driven intestinal inflammation, as shown by the reduction in the histopathological score compared to T cell transfer alone (**Fig. 4B**). Notably, animals receiving the co-transfer of St6-KO B cells exhibited higher histopathological scores than those receiving WT B cells (**Fig. 4B**). Moreover, mice co-transferred with St6-KO B cells showed a lower proportion of IgA^+^ PCs and B cells (**Fig. 4C,D**), with no significant differences in the T cell compartment (**Fig. 4E**). Even though the proportion of total myeloid cells was not altered (**Fig. 4F**), mice that received St6-KO B cells showed a trend toward a higher proportion of neutrophils (**Fig. 4G**). Moreover, a significant decrease in serum IgA was observed in animals co-transferred with St6-KO B cells compared to those receiving WT B cells at weeks 7 and 8 (**Fig. 4H**), with a trend towards lower IgA-coated bacteria in feces (**Fig. 4I,J**), in line with the decrease in IgA^+^ PCs. Thus, selective *St6gal1* deficiency in B cells aggravated the inflammatory response and reduced the frequency of IgA^+^ PCs in both DSS-colitis and the T cell-driven colitis model.

**Figure 4.**
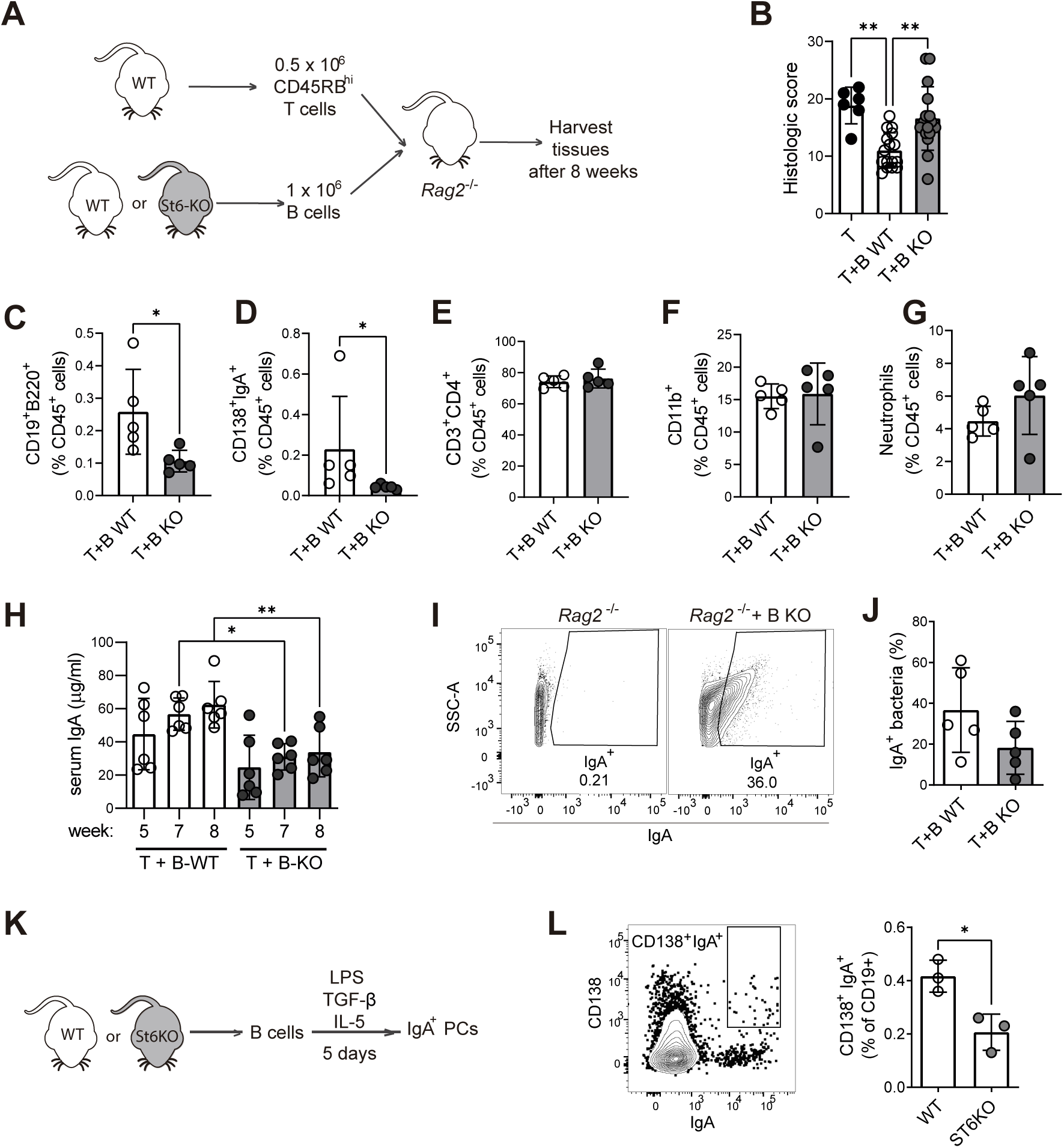
B cells lacking *St6gal1* show impaired amelioration of colitis and differentiation into IgA PCs. **(A-J)** T-cell transfer model of colitis with co-transfer of WT or *St6gal1*^-/-^ B cells. **(A)** Scheme of T cell-adoptive transfer colitis in *Rag2^-/-^*mice that received WT or St6-KO B cells. **(B)** Histopathological assessment of colitis in mice that received only colitogenic T cells (T), T and WT B cells (T+B WT) or T and St6 KO B cells (T+B KO) (mean ± SD; n = 5-16, data are pooled from three independent experiments; one-way ANOVA with Šídák’s multiple comparisons test). **(C-G)** Proportions of LP immune cells: **(C)** CD138^+^IgA^+^ PCs, **(D)** B cells, **(E)** CD3^+^CD4^+^ T cells, **(F)** CD11b^+^ myeloid cells and **(G)** neutrophils (mean ± SD; n = 5, data representative of two or three independent experiments; Student’s t test for all comparisons except IgA^+^ PCs (Mann-Whitney test). **(H)** Quantification of serum IgA by ELISA (mean ± SD; n = 6; data are from one experiment; one-way ANOVA with Šídák’s multiple comparisons test). **(I**) Representative dot plots of IgA binding to fecal bacteria. **Left:** Negative control (*Rag2*^-/-^ mice without B cell transfer). **Right:** *Rag2*^-/-^ mice receiving T and St6KO B cells (T+B KO). **(J)** Percentage of IgA^+^ bacteria (mean ± SD; n = 5, data representative of two independent experiments; Student’s t test). **(K-L)** *In vitro* differentiation of IgA^+^ PCs from WT and St6KO B cells. **(K)** Scheme of the *in vitro* differentiation model. **(L)** Representative dot plot for CD138^+^IgA^+^ PCs and percentage of WT and St6KO CD138^+^IgA^+^ PCs (mean ± SD; n = 3, data representative of three independent experiments; Student’s t test). *p<0.05; **p<0.01; ***p<0.001.

To better understand the functionality of B cells lacking α(2,6)-sialic acid, we analyzed their migratory capacity and differentiation. *St6gal1*^-/-^ B cells showed no significant alterations in migration from the peritoneal cavity to other immune compartments (**Fig. S4A,B**); however, *in vitro* differentiation into IgA^+^ PCs was impaired when compared with WT cells (**Fig. 4K,L**). Thus, lack of α(2,6)-sialic acid on B cells exacerbates the intestinal inflammatory response potentially through a decrease in the frequency of LP IgA^+^ PCs and an increase in neutrophils.

### IgA^+^ PCs and *PIGR*^+^ epithelial cells show an altered transcriptome in UC and DSS-colitis

Considering our previous results, we explored public single-cell transcriptomic profiles of IgA^+^ PCs from biopsies of UC patients compared to non-IBD samples specifically focusing on genes involved in sialic acid biosynthesis, transport, and transfer, as well as those involved in the hexosamine biosynthesis pathway (HBP), as its end product UDP-GlcNAc is a precursor of CMP-sialic acid (**Fig. 5A-C**). Consistent with previous findings [7], IgA^+^ PCs from inflamed tissue showed positive enrichment of several stress-related responses and metabolic alterations as *ATP metabolic process* was one of the most negatively-enriched pathways (**Fig. 5B**). In terms of glycogenes, differential expression analysis (DEA) showed significant upregulation of *ST6GAL1*, *N*-acetylneuraminate synthase (*NANS*, involved in sialic acid biosynthesis) and phosphoglucomutase 3 (*PGM3*, part of the HBP) (**Fig. 5C**). In contrast, a key gene involved in the biosynthesis of the CMP-sialic acid precursor UDP-GlcNAc (UDP-*N*-acetylglucosamine pyrophosphorylase-1, *UAP1*) and several α(2,3)-sialyltransferases (*ST3GAL1*, -*5* and -*6*) were downregulated (**Fig 5C**). Interestingly, neuraminidase-1 (*NEU1*, which removes sialic acid and has been proposed as a negative regulator of B cell activation [35]) was upregulated. Next, we investigated whether these changes in IgA^+^ PCs were also present in DSS-colitis using publicly available data [36]. IgA^+^ PCs from mice with colitis show similar SCPA results, but DEA was limited with non-significant trends (**Fig S5A-C**). These results suggest that *NEU1* and potential decreased bioavailability of UDP-GlcNAc in IgA^+^ PCs may be relevant in inflammation-associated SIgA desialylation.

**Figure 5.**
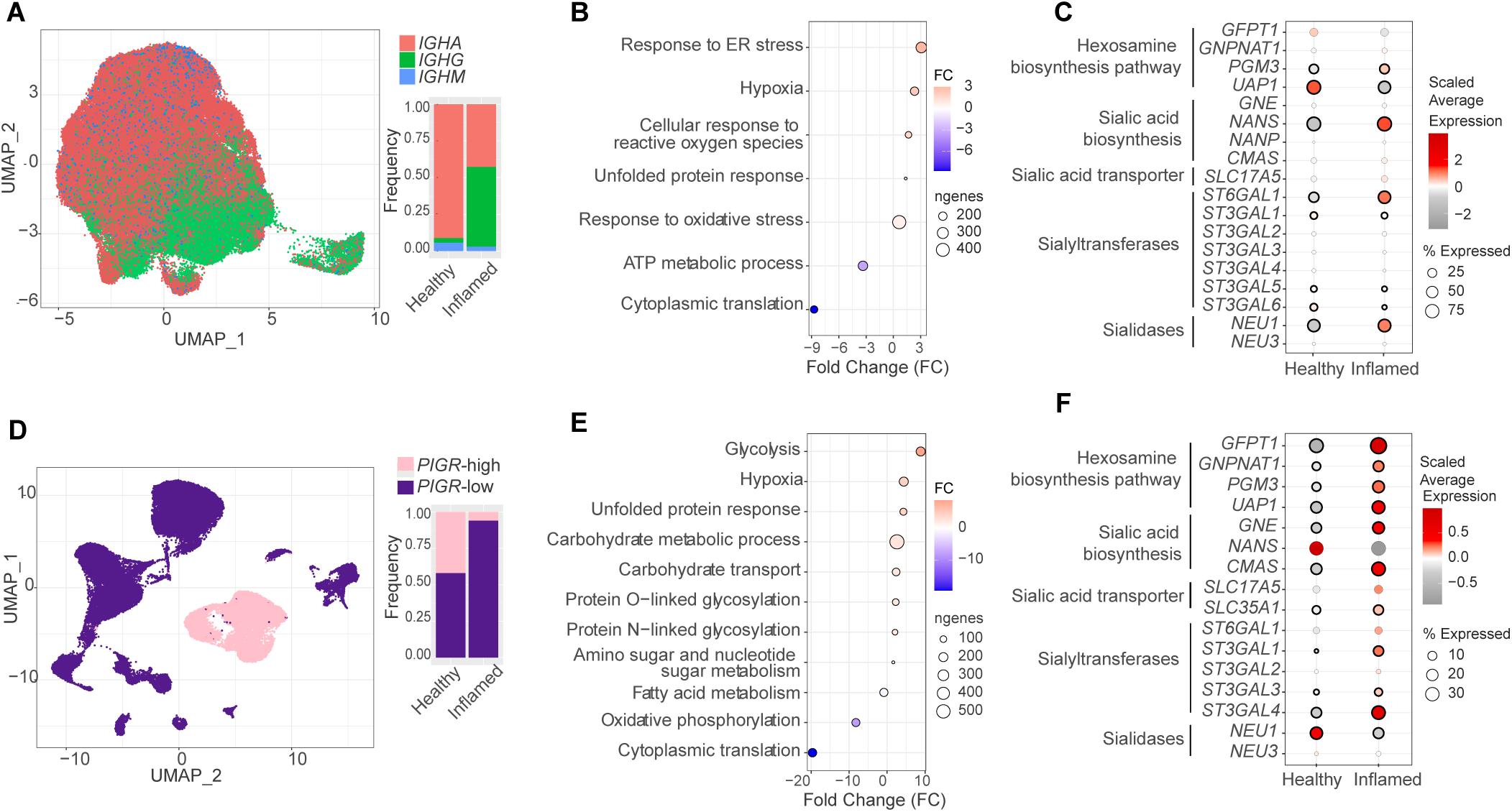
Single-cell transcriptomics analyses of IgA^+^ PCs and *PIGR*-high epithelial cells from UC patients. (**A-C)** Single-cell transcriptomics analysis of IgA^+^ PCs in UC biopsies. **(A)** UMAP shows the distribution of IgA^+^ (*IGHA*), IgG^+^ (*IGHG*) and IgM^+^ (*IGHM*) PCs; barplots show the frequency of each PC isotype relative to total PCs from UC patients (Inflamed) or healthy controls. (**B**) Dotplot representing functional enrichment analyses results obtained using SCPAs. (**C**) Differential expression analysis of IgA^+^ PCs from UC vs healthy controls. Pathways and enzyme families relevant to sialylation status are indicated to the left. (**D-F**) Single-cell transcriptomics analysis of *PIGR*-high epithelial cells in UC. (**D**) UMAP shows the distribution of *PIGR*-high and *PIGR*-low epithelial cells; barplots show the frequency of each group relative to total epithelial cells from UC patients or healthy controls. (**E)** Dotplot representing functional enrichment analyses. (**F)** Differential expression analysis for *PIGR*-high epithelial cells from UC patients (Inflamed) vs healthy controls. For **(B)** and **(E)**, dot color represents the FC of the activity of each pathway comparing inflamed vs healthy control samples, and dot size represents the number of genes composing each pathway. For **(C)** and **(F),** dot color represents the scaled average expression for each gene, and size indicates the proportion of cells expressing each gene in each group, while the black edge represents statistical significance (p<0.05).

Considering that the polymeric immunoglobulin receptor (*PIGR*) is a precursor of the SC in SIgA, we analyzed single-cell data for *PIGR*-high epithelial cells in UC [37] (**Fig 5D-F**). SCPA analysis also showed positively-enriched *unfolded protein response* and *hypoxia*, and a shift in metabolism with decreased oxidative phosphorylation and increased glycolysis (**Fig. 5E**). Notably, both N-and O-glycosylation pathways were positively enriched in these cells (**Fig. 5E**). When specifically analyzing expression of glycosylation-related genes, we observed a significant upregulation of all the HBP enzymes (**Fig. 5F**), as well as Glucosamine-(UDP-*N*-acetyl)-2-epimerase/*N*-acetylmannosamine kinase (*GNE*) and Cytidine monophosphate *N*-acetylneuraminic acid synthetase (*CMAS*), both involved in the biosynthesis of CMP-sialic acid, as well as sialic acid transferases (*ST3GAL1,-3,-4*); in turn, *NEU1* was downregulated (**Fig. 5F**). Thus, transfer and biosynthesis of sialic acid may not be compromised in *PIGR*-high epithelial cells from UC patients, albeit cellular metabolism is altered. In turn, *Pigr*-high epithelial cells in DSS-colitis [36] (**Fig. S5D-F**) showed negative enrichment in several metabolic pathways, as well as O- and N-glycosylation and sialylation (**Fig S5E**). In terms of sialic acid biosynthesis and transfer, these cells showed significant upregulation of *Gne* and downregulation of *Nans,* with alterations in *St3galt3* and *-6* expression (**Fig. S5F**).

## DISCUSSION

Although extensive research into the pathological mechanisms of ulcerative colitis (UC) has deepened our understanding of this disease, a more integrated view of UC is essential for developing more effective therapies. The role of B cells, PCs, and SIgA in intestinal inflammation has recently regained considerable attention [5,6,8,14,17,19]; however, the relevance of their glycosylation and its potential alterations remains poorly understood. Here, using clinical samples and experimental models, we uncover a previously unappreciated role for sialylation in shaping IgA^+^ B-cell responses during intestinal inflammation.

Glycosylation controls diverse immune cell processes, including T cell survival [38], myeloid cell function [39] and B cell activation [40], and influences the transition from healthy to inflamed tissues [41,42]. Particularly, serum immunoglobulin glycosylation has been associated with distinct clinical features of IBD [29,43]. Previous studies have shown that in Crohn’s disease (CD), plasma IgA and fecal SIgA exhibit lower galactosylation and sialylation of N-glycans [19,32], with limited characterization of the glycoprofile in UC, due to less pronounced inflammation-associated alterations [32] or technical limitations [19]. In this study, we show that α(2,6) sialylation is decreased in fecal SIgA of UC patients with active inflammation. Interestingly, this differential glycan profile was not observed during remission, suggesting that it represents a reversible, inflammation-dependent trait. Given that serum IgA in CD exhibits reduced galactosylation and sialylation [32], whereas such alterations were not observed in N-glycans from patients with UC, our findings reveal a potential selective loss of α(2,6)-sialic acid with preserved terminal galactose in fecal SIgA, suggesting that CD and UC may engage distinct pathways to regulate SIgA glycosylation. Moreover, intestinal and systemic IgA pools may undergo aberrant glycosylation through differential regulatory mechanisms. In this context, comparative studies of circulating IgA and fecal SIgA in IBD will be essential to delineate disease-specific glycosylation programs.

SIgA desialylation may result from increased cleavage of sialic acid by host and bacterial neuraminidases during intestinal inflammation [44–46], with α(2,6)-sialidases characterized both in commensal and pathogenic bacteria [47–52]. Further beyond their role in altering mucin glycosylation, neuraminidases could also alter SIgA sialylation as shown in bacterial vaginosis [53,54]. Interestingly, recent reports show that SIgA1 from IBD patients shows increased reactivity to pathobionts [55]. Considering our *in vitro* results showing increased bacterial coating with desialylated SIgA, further studies examining the link between this aberrant glycoprofile and dysbiosis are highly relevant. In this sense, it has also been shown that gut microbiota from *St6gal1* KO mice is enriched in *Helicobacter,* and transfer of their microbiome to a WT recipient induces a mucosal Th17 response [56]. A functionally altered SIgA could trigger or enhance dysbiosis by altered antigen binding and/or transport of immune complexes as previously suggested [19], particularly for abrogated reverse-transcytosis in UC.

Further beyond bacterial neuraminidases, desialylated SIgA could also result from a dysregulated transcriptional profile in IgA^+^ PCs, as well as in epithelial cells, as the source of SC. Even though sialic acid-related pathways in *PIGR*-high epithelial cells are transcriptionally upregulated, the altered metabolism and mitochondrial dysfunction described for these cells in UC [57,58] could limit availability of glycosylation precursors. As epithelial cells are easily accessible to bacterial neuraminidases, their involvement is also plausible.

B cells have a complex, dual role in UC and intestinal inflammation: while specific B cell subsets can exert suppressive and regulatory functions that aid in controlling inflammation [11,12], other B cell populations (such as expanded naïve B cells and IgG^+^ PCs) can promote inflammation and tissue damage [6,9,17,18]. Our results in DSS-colitis reveal a reduction in α(2,6)-sialylation in LP and MLN B cells, as well as in LP IgA⁺ PCs, indicating that the inflammatory milieu during active colitis drives decreased sialylation across the gut-associated B-cell lineage.

Even if genetic deletion of *St6gal1* on B cells is a limited model, our experiments suggest that the protective role of B cells may be dependent on their α(2,6)-sialylation. B cell-specific genetic elimination of *St6gal1* consistently worsened colitis in two experimental models, with a significant decrease of LP CD19^+^B220^+^ and CD138 IgA^+^ cells, as well as increased infiltration of LP neutrophils. Thus, loss of sialylation in the B cell lineage during colitis may be part of a pathogenic circuit that contributes to inflammation. Notably, we found a correlation between loss of α(2,6)-sialic acid and neutrophil infiltration during DSS-colitis in WT mice, as well as an increase in neutrophil infiltration in mice with α(2,6)-sialic acid-deficient B cells. Recent work shows that B cells can induce neutrophil apoptosis through cell-cell contact without relying on classic cell death receptors, thereby promoting reparative macrophage polarization by helping neutrophil phagocytosis [59]. Considering the macrophage/monocyte imbalance observed in intestinal inflammation, and the relevance of Siglec-9 as a mediator of neutrophil activation and apoptosis [60], further work on this subject is warranted.

Our *in vitro* experiments also demonstrated the essential role of α(2,6)-sialylation in the differentiation into IgA^+^ PCs. *St6gal1* KO B cells have already been identified as hyporesponsive, with reduced BCR signaling and proliferation, a trait partly linked to α(2,6)-sialic acid as ligand for CD22, a negative regulator of B cell activation [35,61]. Considering that not only engagement but also CD22 organization can influence B-cell responses, neuraminidases such as Neu1 may also play a crucial role in this process [35].

Interestingly, we observed no differences in intestinal inflammation following the PC-specific genetic ablation of *St6gal1,* which leads us to two non-exclusive hypotheses: a) the exacerbated inflammation observed in experimental models when B cells are devoid of α(2,6)-sialic acid is associated to B cells and not specifically PCs, or b) intestinal inflammation desialylates PCs through mechanisms that are not dependent on *ST6GAL1* expression, thus elimination of this enzyme is redundant. In line with the latter, *ST6GAL1* expression is increased in IgA^+^ PCs in UC and not significantly dysregulated in DSS-colitis. Moreover, the increase in *NEU1* in IgA^+^ PCs from UC patients suggests that a subtle dynamic between these enzymes could regulate cell glycophenotype. Additionally, mislocalization or Golgi redox state alterations (such as those associated to hypoxic microenvironments caused by intestinal inflammation) could affect St6gal1 activity, as previously demonstrated [62]. Finally, we found that in active UC, gut IgA^+^ PCs present a decrease in the expression of *UAP1*, a key enzyme in the HBP and tightly regulated by various factors including cell metabolism [63]. Increasing evidence underscores the differential characteristics of gut B cells and PCs and their metabolic demands, including the singular bioenergetics of gut B cells during differentiation [64,65], the HIF-1α-dependent metabolic reprogramming in IgA class switching [66], and the importance of metabolic fitness for IgA^+^ PCs [67]. Downregulation of *UAP1* in IgA^+^ PC cells from UC patients could compromise UDP-GlcNAc levels; given the crucial role of this enzyme, the upregulation of *PGM3* and *NANS* may not be sufficient to ensure proper sialylation. In DSS-colitis, trends in dysregulation for the HBP and sialylation-related enzymes correlate with our observation that decreased sialylation in IgA^+^ PCs is independent of sialic acid linkage, and with dysregulations observed in active UC. Considering that UDP-GlcNAc is also a key substrate for glycosaminoglycans and O-GlcNAcylation of intracellular proteins [68], the effects of a potential decrease in UDP-GlcNAc levels could also exceed IgA-deficient sialylation.

In summary, our findings uncover a proinflammatory feed-forward circuit in which active colitis induces aberrant glycosylation of SIgA and the gut-associated IgA⁺ B-cell lineage, compromising B-cell function and mucosal homeostasis during intestinal inflammation.

## MATERIALS AND METHODS

### Local patient cohort

All protocols were approved by the Ethics Committee at IBYME-CONICET (Protocol 6848). Samples were obtained with written informed consent. For diagnosis, treatment, and surveillance of UC patients, the European Crohn’s and Colitis Organization (ECCO) guidelines were used. UC patients were classified as mild, moderate or severe based on Truelove & Witts’ criteria for clinical activity [69]. Fecal samples were collected from UC patients and individuals with negative UC diagnosis who attended the Gastroenterology Unit at Hospital Nacional Alejandro Posadas, Argentina (**Table S1**). HIV patients, individuals who received antibiotic therapy 30 days before sample collection, or those with a history of extreme diets (e.g. macrobiotic or vegan), and/or gastrointestinal surgery (including gastrectomy, bariatric surgery, or colostomy) were excluded.

### Human fecal sample collection and purification of SIgA

Participants were instructed on stool sample collection using a standardized written protocol. Approximately 5 g of stool were collected as previously described [70]. For experimentation, a 250 mg aliquot was suspended in PBS, vortexed, and centrifuged (100 *x g*, 15 min); supernatants were diluted in PBS (8,000 *x g,* 5 min) and filtered (0.22 μm Minisart® filter, Sartorius), and IgA was purified using Protein M-agarose (InvivoGen) following the manufacturer’s instructions. Eluates were concentrated (100 kDa-cutoff, Vivaspin, GE Healthcare) and SIgA was quantified by MicroBCA kit (Pierce).

### Mouse strains and *in vivo* experiments

Mice were maintained at the animal facility of IBYME-CONICET under a 12-h light/12-h dark regime. All mice used in this study were on a C57BL/6 background: C57BL/6 WT, *Rag2^−/−^* (B6.Cg-Rag2^tm1.1Cgn^/J, stock no. 008449), *Cd19^cre^* (B6.129P2(C)-Cd19^tm1(cre)Cgn^/J, stock no. 006785), Aicda^cre^ (B6.129P2-Aicda^tm1(cre)Mnz^/J, stock no. 007770) and *St6gal1^f/f^* (B6.129-*St6gal1*^tm2Jxm^/J, stock no. 006901) mice were purchased from the Jackson laboratory. C57BL/6 *St6gal1^−/−^* mice were kindly provided by J. Paulson (La Jolla, CA, USA). *Cd19*^cre^ and *St6gal1*^f/f^ mice were crossed to obtain *Cd19*^cre^ *St6gal1*^f/f^ (B-St6 KO) and *St6gal1*^f/f^ (St6 flox) control mice. *Aicda*^cre^ and *St6gal1*^f/f^ mice were crossed to obtain *Aicda*^cre^ *St6gal1*^f/f^ (PC-St6 KO) and *St6gal1*^f/f^ (St6 flox) mice. The Institutional Committee for the Care and Use of Laboratory Animals (IBYME) approved all protocols (Protocols 013/2023 and 018/2025). Sample sizes were based on variability observed in pilot studies and previous DSS colitis experiments in our laboratory. To standardize housing conditions, cages were kept on the same rack level. In accordance with ethical guidelines, posture, fur condition, stool consistency, rectal bleeding, and body weight were monitored three times a week. The humane endpoint in colitis models was defined as more than 20% body weight loss; no mice reached this endpoint. In experiments involving different genotypes, the experimenter remained blinded to group allocation. Histopathological analyses were conducted blinded.

### DSS-induced colitis

Colitis was induced as previously described [22]. Please refer to **Supplementary Methods** for details.

### Purification of murine luminal SIgA

Intestines of mice were removed, and the luminal contents collected and stored at -80°C. Samples were homogenized in PBS supplemented with Protease Inhibitor Cocktail (Merck) and centrifuged (15,000 *x g*, 15 min, 4°C). Supernatants were collected and filtered (0.22 μm Minisart®, Sartorius) before using Protein L-agarose (InvivoGen) for immunoglobulin purification. Eluates were concentrated (100 kDa-cutoff Vivaspin 6 concentrator, GE Healthcare) and IgA was determined by ELISA (eBioscience).

### Western and lectin blots

Human fecal SIgA (3 μg) was denatured with 4x Laemmli Sample Buffer (Bio-Rad) supplemented with a 710 mM β-mercaptoethanol solution and run in a 12% acrylamide-bisacrylamide Tris-HCl gel under reducing conditions. In turn, 3-5 μg of murine SIgA was run in a 6% acrylamide-bisacrylamide Tris-HCl gel under denaturing, non-reducing conditions to separate IgA from other immunoglobulins. Human SIgA (Sigma) was used as control. Gels were transferred to PVDF membranes (Roche) (250 mA; 3 h) which were then washed and blocked with PBS-5% BSA overnight (4 °C). For Western blotting, biotinylated anti-human IgA antibody (Southern Biotech) or goat anti-murine IgA antibody (Abcam) (2 h; RT) were used. For lectin blotting, membranes were incubated with biotinylated *Sambucus Nigra Agglutinin* (SNA, Vector Labs) or *Erythrina cristagalli* lectin (ECL, Vector Labs) (1 h; RT). Membranes were washed and incubated with streptavidin-HRP (R&D Systems) (30 min; RT) or swine anti-goat IgG-HRP secondary antibody (Southern Biotech) (1 h; RT) and detected using ECL Prime (Amersham, Cytiva) by G-Box chemiluminescence system (Syngene) using GeneSnap software. Protein bands were analyzed with ImageJ software, pixel density was measured and reported as arbitrary units (AU).

### Adoptive transfer of CD4^+^CD45RB^high^ T cells to *Rag2*^-/-^ mice

CD4^+^CD45RB^high^ T cells were isolated from spleens of 10- to 12 week-old female C57BL/6 WT mice as previously described [22]. Total splenic WT or *St6gal1*^-/-^ B lymphocytes were isolated using the Pan B isolation Kit II (Miltenyi Biotec). *Rag2^-/-^*male or female mice (10 to 12 weeks old) were injected intraperitoneally with 0.5 x 10^6^ CD4^+^CD45RB^high^ T cells (WT) for colitis induction, alone or co-transferred with 1.10^6^ WT or *St6gal1*^-/-^ B cells. Mice were randomly assigned to each experimental group using GraphPad QuickCalcs Randomizer, GraphPad Software, California, USA. Available at: https://www.graphpad.com/quickcalcs/randomize1/.

Body weight was measured weekly and normalized at week 0. Animals were euthanized by cervical dislocation at week 8. Histopathological scoring was performed as previously described [22].

### Indirect Immunofluorescence

Immunofluorescence on paraffin-embedded sections was performed following standard protocols for deparaffinization, rehydration, blocking, and indirect staining. For details, please see **Supplementary Methods**.

### Tissue preparation and flow cytometry

After euthanasia, mesenteric lymph nodes (MLN) and colon were removed and processed as previously described [22]. The only alteration is that colon samples were digested (37°C; 45 min; 200 rpm) in PBS (FBS 5% v/v, Sigma) using Collagenase D (1.5 mg/ml, Roche) and DNAse I (20 U/ml, Thermo Fisher Scientific), disaggregated with gentle MACS^TM^ Dissociator and filtered (70 µm). Mouse single-cell suspensions were stained with Zombie Aqua, Green or Violet according to manufacturer’s instructions (Biolegend), washed, and incubated with TruStain Fc (anti-mouse CD16/32) (Biolegend) (10 min, 4 °C) followed by specific antibodies (20 min, 4 °C). For intracytoplasmic IgA, cells were fixed with 2% PFA (30 min, RT), permeabilized and stained with Perm/Wash buffer (BD Bioscience) (30 min, RT). Details on the used antibodies can be found in **Table S2**. Glycophenotyping was performed as previously described [38]. For details, please refer to the corresponding section in **Supplementary Methods**.

### Cytokines and IgA determination

Blood was extracted by submandibular collection or cardiac puncture without anticoagulants in anesthetized mice (ketamine and xylazine). Serum was collected after centrifugation (10 min, 1000 *x g*). Fecal samples were collected and suspended (10 μl PBS/mg) and then homogenized and centrifuged at 12000 *x g* (10 min, 4°C). Colon protein extracts were obtained as previously described [22], and protein concentration was measured with Micro BCA kit (Pierce). IgA (eBioscience) and Lipocalin-2 (RyD) were measured by ELISA. Cytokines were measured using the Legendplex Mouse Inflammation Panel (Biolegend).

### *In vitro* IgA^+^ plasma cell differentiation

Protocol was adapted from [71]; briefly, splenic B cells from WT and *St6gal1^-/-^* animals were cultured with TGF-β (4 ng/ml), IL-5 (10 ng/ml) and LPS (15 μg/ml) for 5 days.

### Functional evaluation of B cells and SIgA

Methods for B-cell homing assays and SIgA binding to bacteria are described in **Supplementary Methods.**

### Single-cell data analysis

Data for cells in DSS-induced colitis was obtained from the GEO database (GSE264408) [36]. In UC, data for PCs was obtained from the Single Cell Portal database (SCP1690) [7], while for epithelial cells the GEO database GSE21469 [37] was used (**Table S3**). Single-cell RNA-seq data was processed as described in Seurat package vignettes [72]. PCs in the DSS model of colitis dataset [36] were classified by using isotype scores calculated using the *AddModuleScore* function from Seurat package (IgA score: *Igha*; IgG score: *Igh1*+*Igh3*; IgM: *Ighm*; IgD: *Ighd*) and defining score-specific cutoffs. Cells that only met the cutoff for one of the scores were annotated, while the others remained “unclassified”. Finally, we calculated a cell cycle score [8] and removed cells expressing high levels of cell cycle genes from the analysis. In UC, PCs were annotated into IgA^+^, IgG^+^ and IgM^+^ subsets by leveraging single-cell V(D)J-sequencing as described by the authors [7]. Epithelial cells both in DSS colitis and UC were grouped based on high and low *PIGR* expression, taking the median expression as the cutoff [36,37].

### Differential expression and enrichment analysis

Seurat’s function “FindMarkers” was used to compare selected identities with default method (Wilcoxon test). Genes were considered differentially expressed with p_adj_<0.05 cutoff. Enrichment analyses were performed using SCPA (Single-cell Pathway Analysis) package with the *compare_seurat* function [73]. The Broad Institute’s MSigDB database pathways under the Gene Ontology categories, as well as the HALLMARKS collection, and the KEGG and REACTOME databases were used and tested for enrichment of those with less than 500 genes. In DSS-colitis, we used the same pathways annotated for mice.

### Statistical analysis and graphical representation of data

Statistical and graphical analyses were performed using GraphPad Prism version 8 (GraphPad Software Inc) or R software version 4.4.2. A significance level (α, probability of making a type I error) of 0.05 was used in all statistical tests. Normality was evaluated using the Shapiro-Wilk test. If one of the groups did not follow a normal distribution, a non-parametric test was selected.

## ACKNOWLEDGMENTS

This work was supported by grants from Agencia Nacional de Promoción Científica y Tecnológica (ANPCyT), PICT 2015-0564 and 2018-2766 to KVM, as well as the National Academy of Medicine/ Fiorini Foundation grant 2024 to KVM. Additionally, we thank Sales, Williams and Baron Foundations for their support to GAR (Argentina), and Fundación Bigand for institutional support. We finally thank the Ferioli, Ostry, Caraballo and Alfonzo families for generous support.

AJC, DOC, APS, FB, GAR, MAT and KVM are research scientists at CONICET, Argentina. AMC, JPM, ADR, LGM, PAG, MNMC, MM, and VMA were supported by doctoral and/or postdoctoral fellowships from CONICET.

## AUTHOR CONTRIBUTIONS

Investigation: AMC, AJC, JPM, ADR, PAG, MNMC, LGM, MM, VCMA, RMM, SGG, M. May, RS, CC, JC, CM. Formal analysis: AMC, AJC, JPM, PAG. Validation: AMC, AJC, JPM, KVM. Visualization: AMC, AJC, JPM, PAG, MAT, KVM. Data curation: AMC, JPM, APS, FSB, KVM. Conceptualization: AMC, MAT, KVM. Methodology: AMC, AJC, JPM, APS, PAG, DOC, FSB, GAR, MAT, KVM. Resources: ADR, DOC, APS, FSB, GAR, KVM. Supervision: CM, DOC, FSB, MAT, KVM. Funding acquisition: GAR, KVM. Project administration: KVM. Writing – original draft: AMC, JPM, MAT, KVM. Writing – review & editing: All authors.

## SUPPLEMENTARY DATA

Supplementary information containing **Tables S1-S3**, **Figures S1-S5** and **Supplementary methods** are available.

## Abbreviations

SIgA: secretory immunoglobulin A
St6gal1: β galactoside α(2,6)-sialyltransferase 1
PCs: plasma cells
BCs: B cells

