## Supplementary figures and materials for "An Altered Glycome Shapes IgA B-Cell Responses and Gut Immunity During Intestinal Inflammation"

**Index**

**Table S1.** Clinical description of the local cohort of UC patients.

**Table S2**. Antibodies used in this study.

**Table S3.** Single cell RNA-seq datasets used in this study.

**Fig. S1**. Changes in human SIgA binding to fecal bacteria after enzymatic desialylation.

**Fig. S2.** Analysis of the immune infiltrate and glycophenotyping of B cells and IgA$^+$ plasma cells in DSS-induced colitis.

**Fig. S3.** St6Gal1 deficiency specifically on B cells exacerbates colitis in the DSS model.

**Fig. S4.** Effects of α(2,6) sialylation on B cell migration and SIgA binding to fecal bacteria.

**Figure S5.** Single-cell transcriptomics analyses of IgA$^+$ PCs and *Pigr*-high epithelial cells from DSS-induced colitis.

**Supplementary Methods.**

**Table S1.** Clinical description of the local cohort of patients. HI, healthy individuals (non-IBD); UC, ulcerative colitis.

| | UC patients | | | HI |
|---|---|---|---|---|
| | UC-inactive[a] | UC-active[a] | | |
| | Clinical remission | Mild | Moderate | |
| Participants in this study | 3 | 3 | 2 | 4 |
| Sex (F/M) | 1/2 | 1/2 | 1/1 | 2/2 |
| Median age (years) (IQR 25%-75%) | 35 (34.5 – 42.5) | 36 (30.5 – 46.5) | 48.5 (42.75 – 54.25) | 44.5 (38.25 – 49.5) |
| Median disease duration (years) (IQR 25%-75%) | 10 (9 – 12) | 9 (9 – 10.5) | 5 (5 – 5) | - |
| Active smokers | 1 | 0 | 0 | 2 |
| **Location UC** | | | | |
| E1 - proctitis | 1 | 0 | 1 | |
| E2 - left-sided | 0 | 1 | 0 | |
| E3 - pancolitis | 2 | 2 | 1 | |
| **Medication** | | | | |
| 5-ASA | 3 | 2 | 2 | |
| Steroids | 0 | 0 | 0 | |
| 6-MP / AZA | 0 | 0 | 0 | |
| 5-ASA / steroids | 0 | 0 | 0 | |
| 5-ASA / AZA / 6-MP | 0 | 0 | 0 | |
| 5-ASA / AZA/ steroids | 0 | 1 | 0 | |
| 5-ASA / ATB | 0 | 0 | 0 | |
| AZA / ATB | 0 | 0 | 0 | |
| Steroids / ATB | 0 | 0 | 0 | |
| **No medication** | 0 | 0 | 0 | |

IQR, interquartile range; 5-ASA, 5-aminosalicylic acid (mesalamine); 6-MP, 6-mercaptopurine; AZA, azathioprine; ATB, antibiotics; upper GIT, upper gastrointestinal tract; 0, non-existent; -, not applicable.

[a] Activity based on Truelove-Witts criteria.

**Table S2. Antibodies used in this study.**

| Antibody | Clone | Source | Catalog |
|---|---|---|---|
| FITC anti-IgA mouse | C10-3 | BD Biosciences | 559354 |
| PE anti-IgA mouse | ma-6e1 | eBioscience | 12-4204-82 |
| PE anti-IgA mouse | 11-44-2 | SouthernBiotech | 1165-09 |
| APC anti-IgA mouse | 11-44-2 | SouthernBiotech | 1040-31 |
| anti-IgA mouse | polyclonal | Abcam | ab97231 |
| BV510 anti-CD45 mouse | 30-F11 | Biolegend | 103137 |
| BV785 anti-CD45 mouse | 30-F11 | Biolegend | 103149 |
| APC-Fire 750 anti-CD19 mouse | 6D5 | Biolegend | 115558 |
| APC anti-B220 mouse | RA3-6B2 | Biolegend | 103212 |
| BV711 Anti- B220 mouse | RA3-6B2 | Biolegend | 103255 |
| FITC Rat anti-CD3 mouse | 145-2C11 | Biolegend | 100306 |
| AF647 anti-CD3 mouse | 17A2 | Biolegend | 100209 |
| APC-Cy7 anti-CD4 mouse | RM4-5 | Biolegend | 100526 |
| AF700 anti-CD4 mouse | 17A2 | Biolegend | 100216 |
| PerCP-Cy5.5 anti- CD8 mouse | 53-6.7 | Biolegend | 100734 |
| FITC anti- CD8 mouse | 53-6.7 | Biolegend | 100706 |
| APC-Cy7 anti-Ly6C mouse | HK1.4 | Biolegend | 128026 |
| APC-Cy7 anti-Ly6C mouse | HK1.4 | Biolegend | 128049 |
| PE-Cy7 anti-Ly6G mouse | 1A8 | Biolegend | 127618 |
| BV605 anti-CD11b mouse/human | M1/70 | Biolegend | 101257 |
| BV421 anti-CD138 mouse | 281-2 | Biolegend | 142523 |
| BV785 anti-CD138 mouse | 281-2 | Biolegend | 142534 |
| PerCP-Cy5.5 anti-I-A/I-E mouse | M5/114.15.2 | Biolegend | 107625 |
| AF488 anti-I-A/I-E mouse | M5/114.15.2 | Biolegend | 107616 |
| APC anti-CD170 mouse (Siglec-F) | S17007L | Biolegend | 155508 |
| FITC anti-CD170 mouse (Siglec-F) | S17007L | Biolegend | 155504 |
| APC/Cyanine7 anti-CD11c mouse | N418 | Biolegend | 117324 |
| FITC anti-CD45RB mouse | C363-16A | Biolegend | 103306 |
| BV421 anti-CD64 (FcγRI) mouse | X54-5/7.1 | Biolegend | 139309 |
| Biotin Goat anti-IgA human | polyclonal | SouthernBiotech | 2050-08 |

**Table S3. Single cell RNA-seq datasets used in this study.**

| Accession number | Cell type analyzed | Subjects | Subset of samples used in this study | Comparison | Number of cells | Number of DEGs | Reference |
|---|---|---|---|---|---|---|---|
| GSE264408 | Mouse plasma cells | 3 healthy controls<br><br>3 acute colitis<br><br>4 chronic colitis | 3 Control<br><br>4 Chronic Colitis | IgA$^{+}$ cells of chronic colitis samples vs healthy controls | Healthy: 354<br><br>Chronic colitis: 751 | Up: 50<br><br>Down: 48<br><br>NS: 7466 | Hong et al., 2024 [36] |
| | Mouse epithelial cells | | | *Pigr*-high cells of chronic colitis samples vs healthy controls | Healthy: 957<br><br>Chronic colitis: 238 | Up: 182<br><br>Down: 284<br><br>NS: 10451 | |
| SCP1690 | Human plasma cells | 8 healthy controls<br><br>8 UC active<br><br>5 UC remission | 8 Control<br><br>8 UC active - Inflamed tissue | IgA$^{+}$ cells of inflamed tissue vs healthy controls | Inflamed tissue: 26874<br><br>Healthy control: 42969 | Up:3401<br><br>Down: 2749<br><br>NS: 4340 | Scheid et al, 2023 [7] |
| GSE214695 | Human epithelial cells | 6 healthy controls<br><br>6 UC | 6 Control<br><br>6 UC | *PIGR*-high cells of inflamed tissue vs control | Healthy control: 6299<br><br>Inflamed tissue: 885 | Up: 3291<br><br>Down: 6253<br><br>NS: 3602 | Garrido-Trigo et al., 2023 [37] |

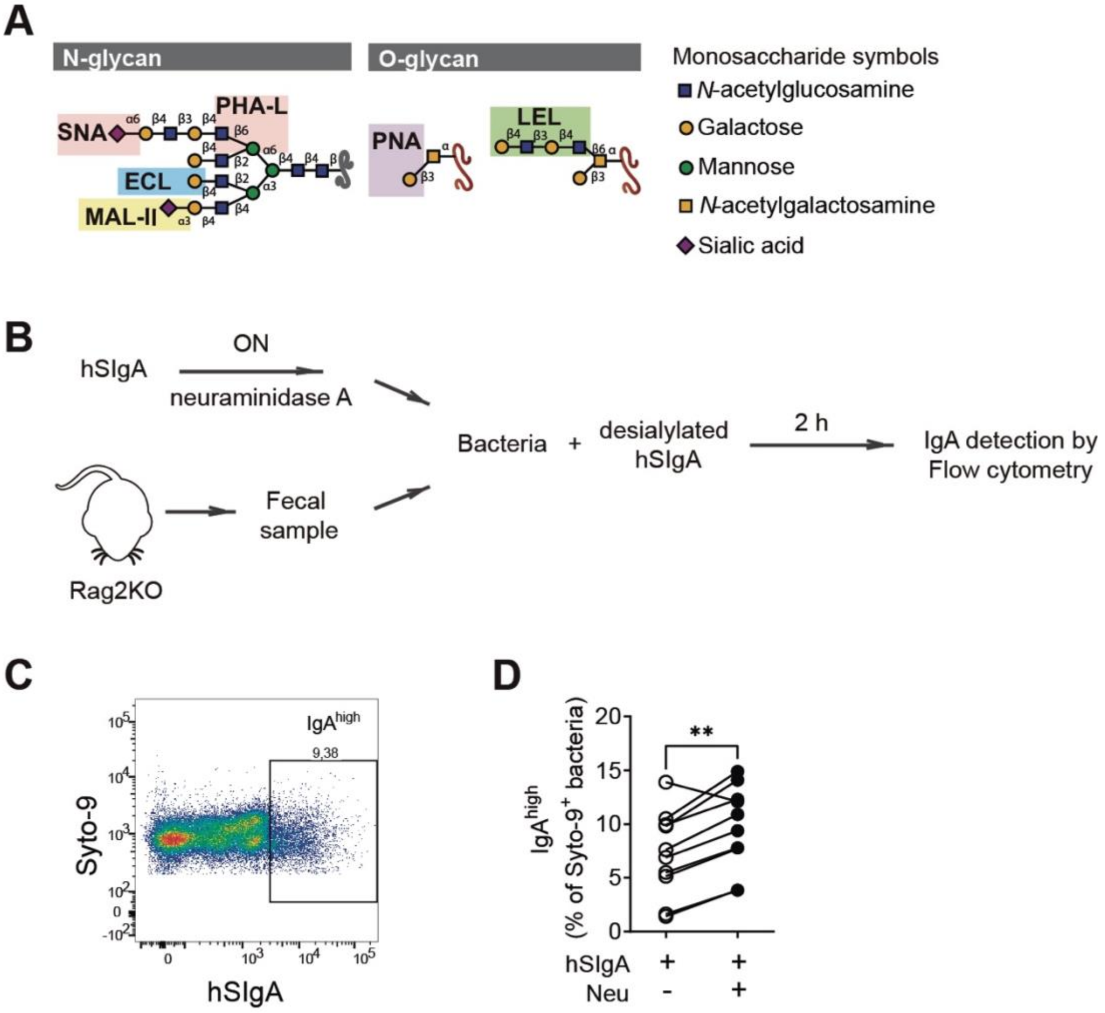


**Fig. S1. Changes in human SIgA binding to fecal bacteria after enzymatic desialylation.** (**A**) Schematic representation of lectin recognition patterns. Glycans are depicted following the guidelines from the Symbol Nomenclature for Glycans (SNFG Discussion Group, Neelamegham et al, 2019). **(B)** Experimental scheme for the evaluation of hSIgA binding to bacteria after enzymatic desialylation, using immunoglobulin-deficient *Rag2*$^{-/-}$ mice as fecal donors. **(C)** Dot plot showing gating strategy for hSIgA$^{high}$ bacteria. **(D)** Binding of hSIgA treated or not with neuraminidase A (Neu) to fecal bacteria (each pair of dots represents the sample from a mouse, n = 10, data are pooled from two independent experiments; paired t-test). **$p<0.01$.

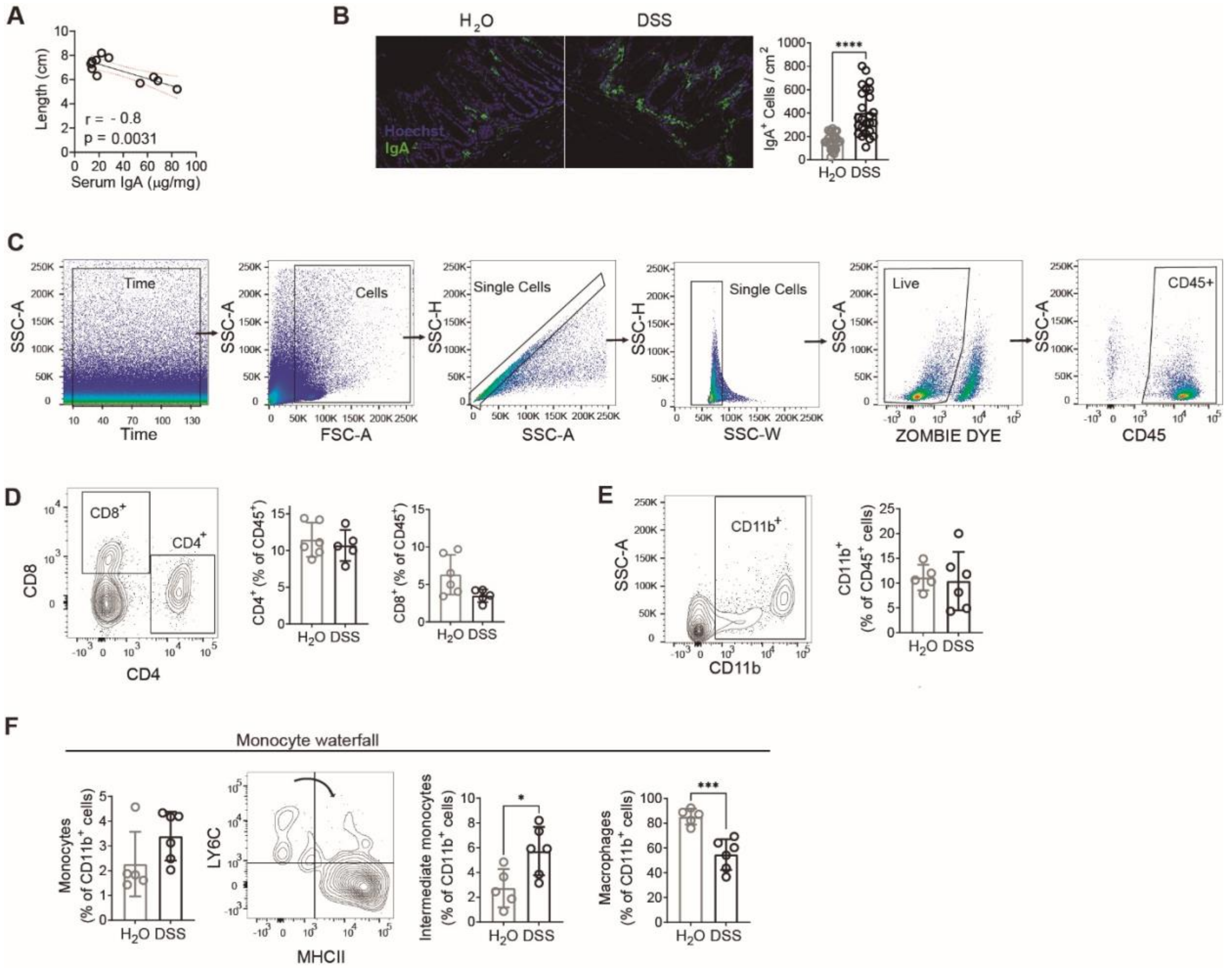


**Fig. S2. Analysis of the immune infiltrate and glycophenotyping of B cells and IgA$^+$ plasma cells in DSS-induced colitis. (A)** Correlation between colonic length and serum IgA in control and DSS-treated mice. Solid black line indicates the best fit curve, dotted red lines indicate 95% trust intervals. Pearson´s p-value (p) and correlation coefficient (r) are shown, n = 11. **(B)** Representative confocal microscopy with IgA in the colon of DSS-treated and control mice (left) and quantification of IgA$^+$ cells/cm$^2$ (right). IgA staining in green (ALEXA 488) and nuclei (DAPI) in blue (mean ± SD, n = 25 random locations, data pooled from two independent experiments; Welch's t-test). **(C)** Gating strategy for colonic CD45$^+$ immune cells. **(D)** Representative flow cytometry staining for T cells and proportions of CD4$^+$ and CD8$^+$ cells in lamina propria (LP) from control and DSS-treated mice. **(E)** Proportions and representative dot plots of total CD11b$^+$ myeloid cells. **(F)** Colonic monocyte/macrophage subset proportions and representative dot plot. Monocytes are selected as Ly6C$^{high}$MHCII$^{low}$, Intermediate monocytes as Ly6C$^{high}$MHCII$^{high}$ and macrophages as Ly6C$^{low}$MHCII$^{high}$. (**D,E,F**) data is shown as mean ± SD; n = 5-6, Student's t test for all comparisons except monocytes (Mann Whitney test). *$p<0.05$, **$p<0.01$, ***$p<0.001$.

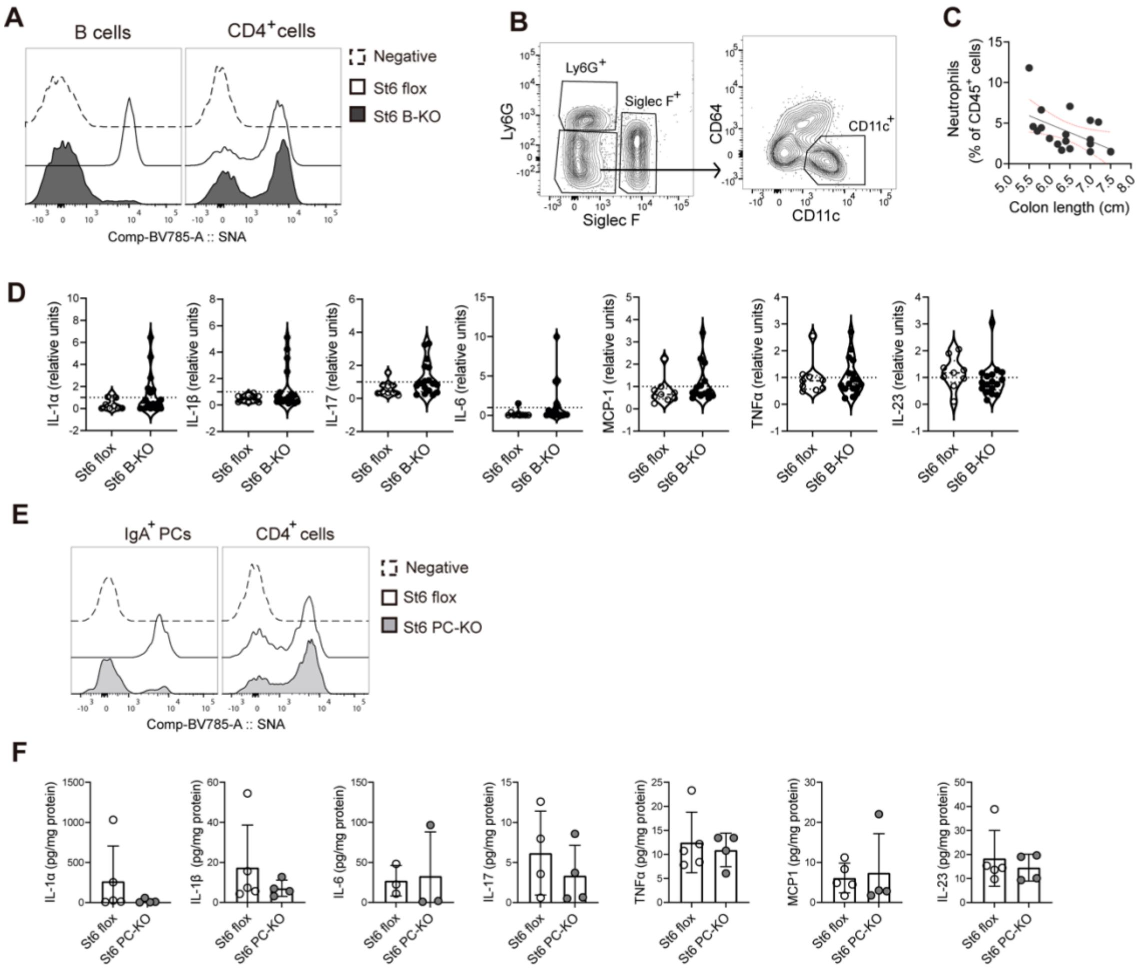


**Fig. S3. St6Gal1 deficiency specifically on B cells exacerbates colitis in the DSS model. (A-C)** DSS-induced colitis in *Cd19*$^{Cre}$ *St6gal1*$^{f/f}$ (St6 B-KO) and *St6gal1*$^{f/f}$ (St6 flox) littermate mice. **(A)** Representative histogram showing the binding of SNA to LP B cells and CD4$^{+}$ T cells in St6 flox and St6 B-KO mice with negative control of SNA. **(B)** Gating strategy for subsets of CD11b$^{+}$ cells: neutrophils selected as Ly6G$^{+}$ cells, eosinophils as Siglec F$^{+}$ cells and dendritic cells as CD11c$^{+}$ cells. **(C)** Correlation between LP neutrophils and colon length in control and DSS-treated mice. Solid black line indicates the best fit curve, dotted red lines indicate 95% trust intervals. Spearman´s P-value (p) and correlation coefficient (r) are shown, n = 19). **(D)** Violin plots depicting relative levels of colon cytokines measured by bead array (n = 9-17, data are pooled from two independent experiments, levels of cytokines within each experiment were relativized by dividing with corresponding experiment mean; Mann Whitney test). **(E,F)** DSS-induced colitis in *Aicda*$^{Cre}$ *St6gal1*$^{f/f}$ (St6 PC-KO) and *St6gal1*$^{f/f}$ (St6 flox) littermate mice. **(E)** Histogram showing the binding of SNA to LP IgA$^{+}$ PCs and CD4$^{+}$ T cells in St6flox and St6 PC-KO mice with negative control of SNA. **(F)** Levels of colon cytokines measured by bead array (mean ± SD; n = 4-5, data are from one experiment; Mann Whitney test).

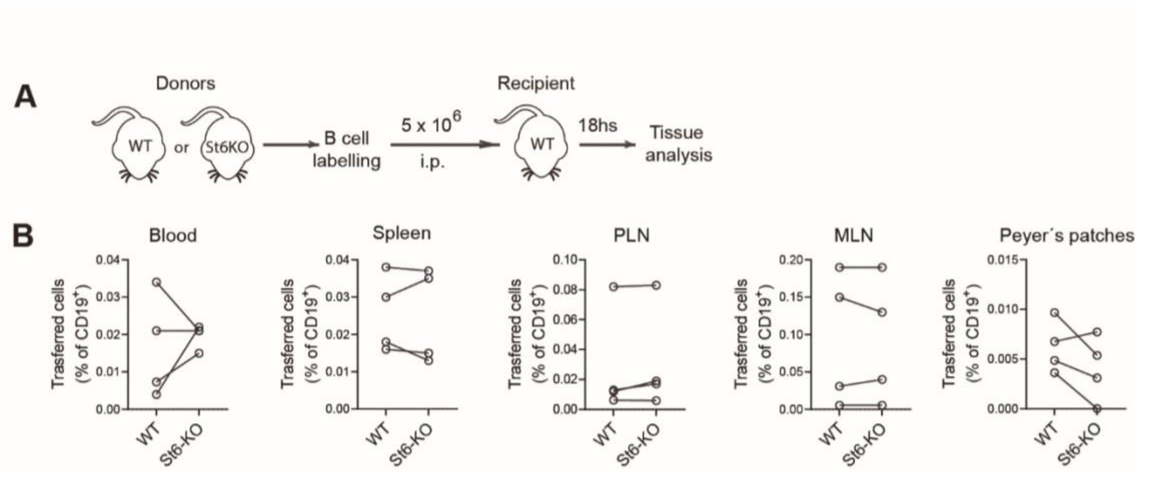


**Fig. S4. Effects of α(2,6) sialylation on B cell migration. (A)** Experimental scheme for the evaluation of B cell migration *in vivo*. **(B)** Percentage of transferred cells (positive for CD19 and their respective labels) in different tissues (each pair of dots represents one mouse; n = 4, data of one experiment; paired t-test). PLN: peripheric lymph nodes; MLN: mesenteric lymph nodes.

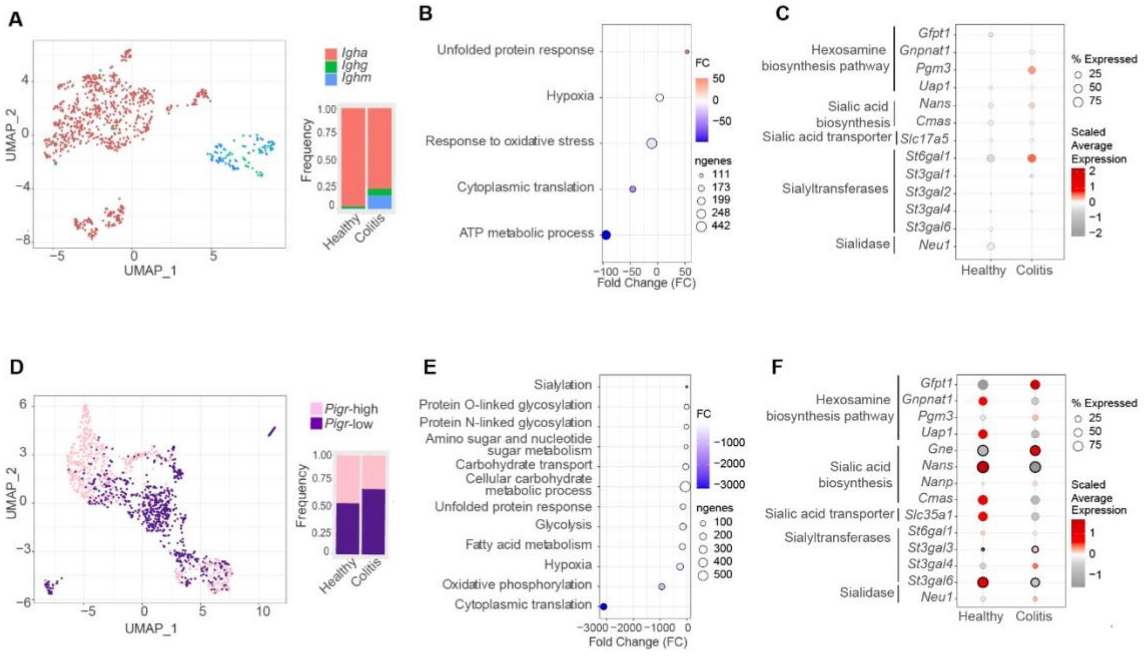


**Figure S5. Single-cell transcriptomics analyses of IgA$^+$ PCs and *Pigr*-high epithelial cells from DSS-induced colitis (Hong *et al.*, 2024, [37]).** (**A-C)** Single-cell transcriptomics analysis of IgA$^+$ PCs in DSS-induced colitis. **(A)** Left: UMAP shows the distribution of IgA$^+$ (*IGHA*), IgG$^+$ (*IGHG*) and IgM$^+$ (*IGHM*) PCs. Right: Barplots show the frequency of each PC isotype relative to total PCs from mice with chronic colitis (colitis) vs healthy controls. (**B**) Differential expression analysis of IgA$^+$ PCs from colitis vs healthy controls. (**C**) Dotplot representing functional enrichment analyses results obtained using *Single-Cell Pathway Analysis* (SCPAs). Pathways and enzyme families relevant to sialylation status are indicated to the left. (**D-F**) Single-cell transcriptomics analysis of *Pigr*-high epithelial cells in DSS-induced colitis. (**D**) Left: UMAP shows the distribution of *Pigr*-high and *Pigr*-low epithelial cells. Right: Barplots show the frequency of each group relative to total epithelial cells from colitis vs healthy control mice. (**E**) Differential expression analysis for *Pigr*-high epithelial cells from colitis vs healthy control mice. (**F**) Dot plot representing functional enrichment analyses. For **(B)** and **(E)**, the color of the dot represents the FC of the activity of each pathway comparing colitis vs healthy control samples, the size of the dot represents the number of genes composing each pathway. For **(C)** and **(F)** **t**he color of the dots represents the scaled average expression for each gene, the size indicates the proportion of cells expressing each gene in each group, black edge on the dots represents statistical significance ($p<0.05$).

### Supplementary Methods

#### DSS colitis

Briefly, 8- to 12-week-old female and male C57BL/6 mice received 2.5% (w/v) DSS (MW 36,000 to 50,000; MP Biomedicals) in drinking water (three cycles of 5 days of treatment followed by 5 days of water). Animals were euthanized on day 30 by cervical dislocation for analysis. In experiments comparing water- and DSS-treated WT animals, investigators were not blinded to group allocation during experimental procedures but were blinded during data analysis.

#### Glycophenotyping by flow cytometry

Briefly, after viability staining cells were incubated with biotinylated lectins (30 min; 4°C) followed by fluorochrome-conjugated streptavidin (Biolegend). The relative median of fluorescence intensity (rMFI) was calculated as (median intensity of cells of interest / median mean intensity of negative control). Data were obtained using a FACSCanto II or LSRFortessa flow cytometer (Becton Dickinson) and analyzed with FlowJo software (version 10.7.1; FlowJo, LLC).

Lectins used for glycophenotyping include phytohemagglutinin-L (L-PHA; recognizing β(1,6) branched complex N-glycans), *Sambucus nigra agglutinin* (SNA, with specificity for α(2,6) sialic acid), *Maackia amurensis* lectin II (MAL-II, recognizing α(2,3) sialic acid), peanut agglutinin (PNA; with specificity for asialo-core 1 O-glycans), and *Lycopersicon esculentum* lectin (LEL, recognizing poly-LacNAc residues).

#### Indirect Immunofluorescence

Paraffin-embedded sections were cleaned and hydrated by incubation at 37°C for 1 h and immersion in xylene (1h), followed by ethanol 100% and decreasing dilutions of this solvent (96%, 80%, 50%) in water. Autofluorescence was removed using ammonium chloride 50 mM (1h, RT). Blocking was performed with 5% BSA in PBS. Anti-IgA mAb (Abcam) was incubated (1:300 in PBS, BSA 1%) overnight at 4°C. After washing, a secondary anti-goat ALEXA 488 was added (1 h, RT) and then washed three times with PBS. Finally, nuclei were stained with Hoescht on Vectashield mounting medium (Vector). Photos were taken from random locations (n=25) and analyzed using ImageJ software; $IgA^+$ cells were counted manually and relativized to area units.

**Homing assays**

Splenic B cells from WT and *St6gal1*$^{-/-}$ mice were incubated (15 min, 37 °C) with 1 µM CellTrace™ Far Red dye (CellTrace™ Far Red Cell Proliferation Kit, Life Technologies) and Tag-it Violet™ Proliferation and Cell Tracking Dye (Biolegend) respectively. After labeling, $5x10^6$ WT cells and *St6gal1*$^{-/-}$ were injected (i.p.) into WT mice, and animals were euthanized after 18 h. Blood, MLN, axillary and inguinal nodes, Peyer's patches, and spleens were removed, tissues were disrupted and cells were stained for cytometry.

**SIgA binding to bacteria**

Feces or intestinal contents were collected from *Rag2*$^{-/-}$ mice, mechanically disintegrated in 100 µl of cold PBS/10 mg, and centrifuged (10 min, 100 *x g*). Approximately 50 µl of the supernatant were re-centrifuged and the resulting pellet (8000 *x g*) was washed twice and incubated for 15 min with the DNA probe SYTO 9 (ThermoFisher). For hSIgA binding, the pellet was incubated (2 h on ice) with human colostrum SIgA (Sigma) that was digested (or not) ON with 3 µl of α(2-3,6,8,9) neuraminidase A (New England BioLabs) following manufacturer´s instructions. Then, bacteria were washed and incubated with biotinylated anti-human IgA (20 min, SouthernBiotech), followed by streptavidin-BV421 (Biolegend). For murine IgA detection, bacteria were labeled with anti-murine IgA-PE (Southern Biotech) (20 min on ice).
